# Clonal expansion accounts for most post-disturbance proliferation of a dominant temperate liana, *Wisteria floribunda*, across a fragmented forest landscape

**DOI:** 10.1101/2025.02.03.636348

**Authors:** Hideki Mori, Takashi Kamijo

**Author notes:** Corresponding author: Hideki Mori.

## Abstract

Large-scale anthropogenic disturbances, including forest fragmentation, are increasing worldwide. Lianas often proliferate after such events, yet the mechanisms underlying these increases— particularly their demographic sources—remain unclear. Here, we quantify the extent to which clonal expansion, rather than seed recruitment, accounts for post-disturbance proliferation of *Wisteria floribunda* (Fabaceae), the dominant liana in the study area. In a 21.6-ha fragmented forest landscape in Japan, we divided the study area into 20 × 20 m grid cells and randomly sampled up to one *W. floribunda* ramet per cell, and genotyped 273 ramets at 10 microsatellite loci to assign ramets to genets and quantify genet areas and their spatial distribution. Genotyping revealed a substantial contribution of extensive clonal reproduction: 94% of ramets were clonal, resolving into 47 genets. These genets collectively occupied 58% of the 21.6-ha study area, and expansion produced extensive genets—the largest covered 9.5 ha. Several genets spanned multiple forest patches; three multi-patch genets accounted for 43% of all ramets, including one spanning seven patches. Multi-patch genets were concentrated within areas covered by forests prior to fragmentation (odds ratio = 6.83; P < 0.001). These observations indicate that clonal expansion accounts for most post-disturbance proliferation, that some genets attain landscape-scale coverage in a fragmented forest, and that part of this dominance likely reflects legacy effects of genets established prior to fragmentation. Explicitly accounting for clonal contributions and genet spatial scale is critical for accurate prediction of disturbance-driven liana dynamics.

## 1. Introduction

In recent decades, deforestation has escalated rapidly, driving widespread forest fragmentation on a global scale and prompting concerns over its profound impacts on biodiversity and ecosystem functioning (Hansen et al., 2013; Liu et al., 2018). Such fragmentation disrupts habitat structure and connectivity, altering ecological processes across diverse taxa (Haddad et al., 2015; Li et al., 2022; Kiesewetter et al., 2023) and ultimately affecting ecosystem functions (Fahrig, 2003; Wilson et al., 2016). Among these impacts, increased liana (woody vine) abundance is particularly noteworthy, with fragmented forests often exhibiting higher liana density and dominance (Campbell et al., 2015; Ladwig and Meiners, 2015). This proliferation is considered a major factor in their global rise relative to trees (Schnitzer and Bongers, 2011; Ngute et al., 2024), yet the specific ecological mechanisms—particularly at the species level—that drive liana expansion in these altered landscapes remain poorly understood.

Post-disturbance liana increases could arise from two non-exclusive processes. On one hand, the high-light environments generated by disturbance favor the recruitment and survival of seed-origin liana seedlings (Laurance et al., 2001; Mori et al., 2020; Yuan, 2024). On the other hand, clonal expansion—vegetative proliferation where a single genetic individual (genet) produces multiple apparent individuals (ramets)—can also be important and may occur at multiple spatial scales: while localized clonality, often inferred from above-ground contiguity, is widely recognized (e.g., Putz, 1984; Alvira et al., 2004; Ledo and Schnitzer, 2014; Schnitzer et al., 2021), some liana species such as *Wisteria floribunda* can form spatially extensive genets in natural forests (Sakai et al., 2002; Mori et al., 2021). These alternatives frame a central uncertainty in forest landscapes fragmented by disturbance—namely, the relative importance of seed recruitment versus large-scale clonal expansion. Because field traits have limitations for inferring genet identity, and underground clonal connections can decay over time, genetic analysis is the most direct way to identify which ramets belong to the same genet and to measure its spatial extent.

In this context, *Wisteria floribunda* (Fabaceae) provides a well-suited system to investigate these strategies. It is widely distributed across temperate forests in East Asia (GBIF Secretariat, 2023) and is regionally dominant in our study area; for example, in a continuous natural forest, *W. floribunda* accounted for 69% of liana stems and 85% of basal area (Mori et al., 2016a; Mori et al., 2025). The species exhibits dual capacities central to this study: in continuous, closed-canopy forests it spreads clonally via stolon extension and can form extensive genets, with ramets separated by up to ∼180 m (Sakai et al., 2002; Mori et al., 2018), and ontogenetic evidence indicates that such lateral clonal expansion typically initiates after ramets reach the bright forest canopy (Mori et al., 2021). Taken together, previous studies suggest a life history in which seed recruitment occurs primarily in high-light environments, whereas once ramets reach the bright forest canopy, stolon-mediated clonal expansion can extend and persist under shaded, closed canopies. In fragmented forests with extensive edge habitat, where light is abundant and small-diameter trees are common (e.g., Laurance et al., 2001), seed recruitment would therefore also be expected to contribute substantially—motivating our test of the relative contributions of these processes.

Here, we quantify and partition the demographic sources—seed recruitment versus clonal expansion—of post-disturbance proliferation at the landscape scale by resolving genets and quantifying genet areas, coverage, and spatial distribution. We ask two related questions: (i) what proportion of *W. floribunda* ramets are clonal versus seed-derived, and (ii) to what extent clonal expansion contributes to the observed proliferation across a fragmented forest landscape. We also ask whether large genets are disproportionately located within areas that were forest prior to fragmentation, a pattern that would be consistent with legacy effects of genets established prior to fragmentation.

## 2. Materials and Methods

### 2.1. Study sites

The study site comprises multiple forest fragments within a 21.6-ha area on the campus of the University of Tsukuba, Ibaraki Prefecture, Japan (36°06′40″N, 140°06′17″E; Figs. 1, S1). To reconstruct forest-cover history, we used aerial photographs (1959–2008; Geospatial Information Authority of Japan) and recent satellite imagery (Copernicus Sentinel; European Space Agency). We mapped forest as areas where tree crowns covered the majority of the area and applied this criterion consistently to the available imagery to identify forest cover for selected years (Fig. S2). Under this criterion, forest area in 1959 was 15.1 ha (∼70% of the 21.6-ha study area; Fig. 2). Forest patches were delineated from aerial photographs as contiguous forest units bounded by roads and other anthropogenic discontinuities where canopy connectivity was absent (Figs. 1, S3).

**Figure 1.**
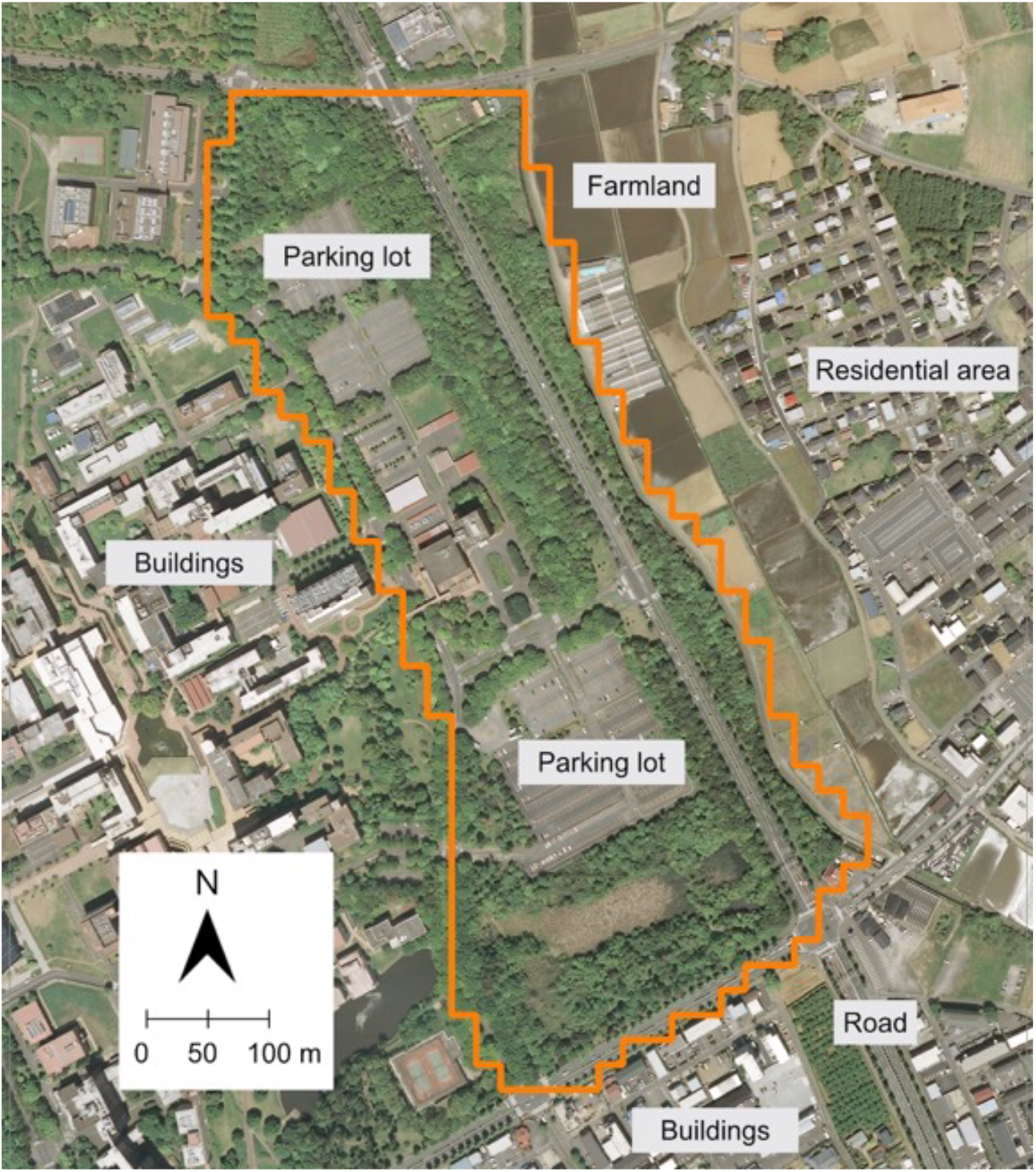
Aerial photograph showing the study area (outlined in orange), which is embedded within an urban landscape. The surrounding area includes buildings, parking lots, roads, and other infrastructure, representing a typical case of anthropogenic forest fragmentation. The photograph was taken in 2008 by the Geospatial Information Authority of Japan.

**Figure 2.**
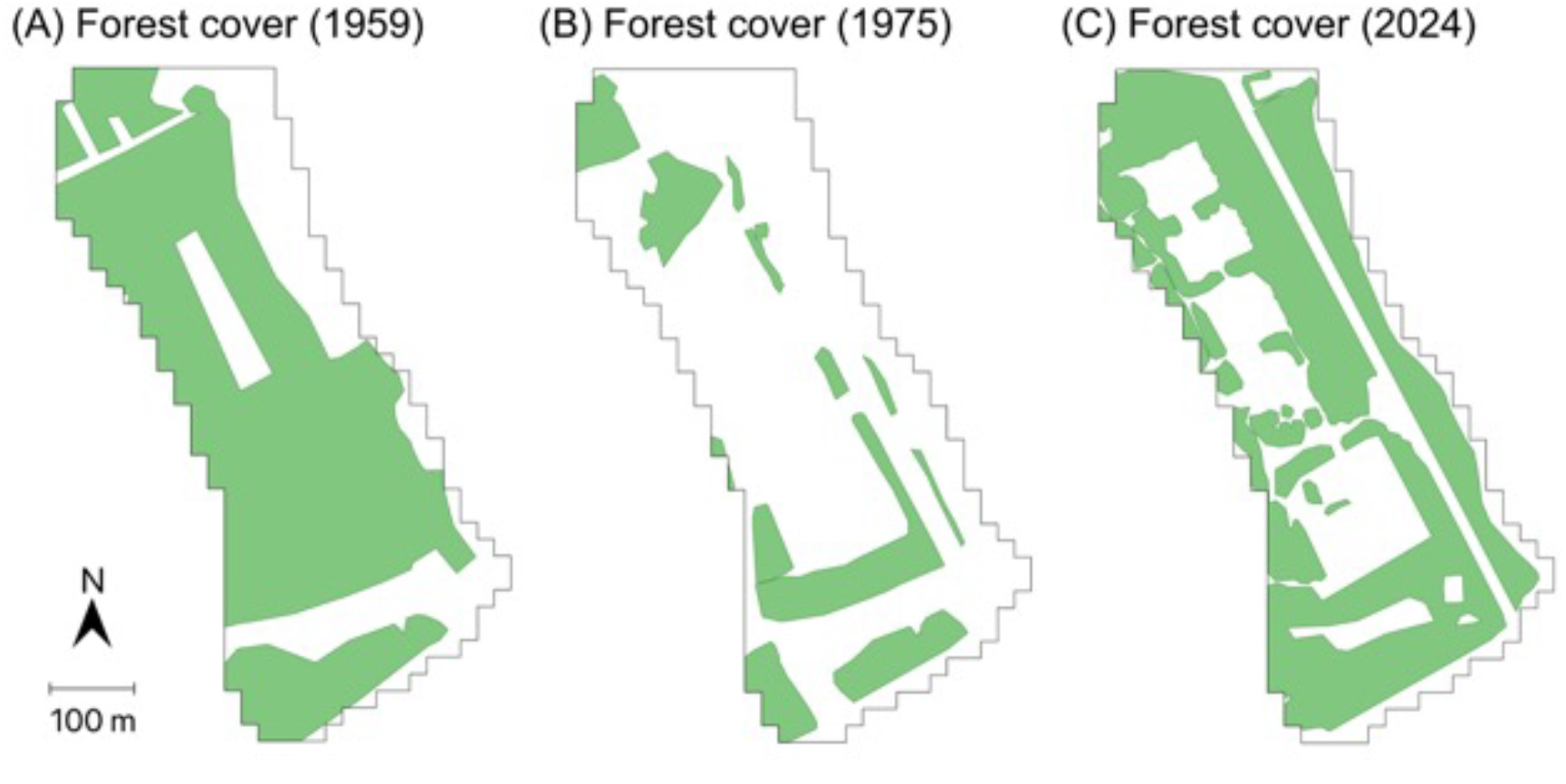
Temporal changes in forest cover at the study site. Panels show (A) forest cover in 1959, when largely continuous forest cover was present before major development; (B) forest cover in the 1970s, showing the minimum forest area after widespread clearing; (C) forest cover in 2024, reflecting partial recovery through afforestation and succession.

Over the subsequent decades, large-scale development in the 1960s–1980s fragmented the forests into multiple smaller patches and reduced total forest area (Figs. 2, S2). By 1975, forest area had declined to 4.6 ha (about 30% of the 1959 forest area). Forest area then increased through secondary succession and afforestation, reaching 12.5 ha by 1998; forest area and patch number have been broadly stable since the late 1990s. At present, areas with frequent human access— primarily along roads—are subject to routine mowing two to three times per year between May and November, whereas most other forested areas are unmanaged (Fig. S4).

### 2.2 Study species

*Wisteria floribunda* (Fabaceae) is a temperate deciduous liana (Fig. S5) (Ohashi, 1989; Sakai and Suzuki, 1999). This species is classified as a stem twiner (sensu Putz and Holbrook, 1991). *Wisteria floribunda* is reported to be a dominant species with high abundance in old-growth temperate forests in the same region as the study site (Ibaraki Prefecture, Japan; Suzuki, 2002; Mori et al., 2016a; Mori et al., 2025), making it a representative liana species in the region. This species is characterized by relatively large stems, with individuals reaching a maximum of approximately 40 cm in diameter under natural conditions (Mori et al., 2018). *Wisteria floribunda* reproduces clonally through the extension of stolons (Sakai et al., 2002), which increases the number of ramets. This species is also a major problematic liana in forestry, as it entwines around planted trees, reducing the quality of harvested timber (Suzuki, 1989). Furthermore, many liana species native to East Asia, including *W. floribunda*, are invasive in regions such as North America, where they pose significant ecological challenges, such as competition with native species (Trusty et al., 2007; Leicht-Young and Pavlovic, 2015).

Given these characteristics, focusing on *W. floribunda* offers valuable insights that range from fundamental ecological knowledge to applied perspectives on management and conservation in liana species. As such, *W. floribunda* is well suited as a focal species for investigating the role of clonal reproduction in the proliferation of lianas under forest fragmentation.

### 2.3 Liana Inventory

To estimate liana abundance and community structure within the study site, we conducted a liana census. The study area was divided into 20 × 20 m grid cells (Fig. S6). Because lianas attached to trees (“on-tree” stage) were almost absent in managed areas, we randomly selected fifteen forested grid cells (total area = 0.6 ha) outside managed areas. Within each selected cell, we recorded the species and diameter at breast height (DBH) of all lianas attached to trees (DBH > 0 cm) on host trees (DBH > 1 cm), following Mori et al. (2025) and Gerwing et al. (2006). Species names followed Yonekura and Kajita (2003).

### 2.4 Sampling, DNA extraction, and Genotyping

The study area, divided into 20 × 20 m grid cells as described above, was used for sampling. We randomly sampled up to one *W. floribunda* plant per grid cell. In each cell, we first checked for on-tree ramets (attached to host trees with tree DBH > 1 cm); if one or more were present, we randomly sampled one. If no on-tree ramet was present, we instead randomly sampled one pre-climbing (“on-floor”) individual (a plant not yet attached to a host). This substitution was used to maintain spatial coverage and avoid bias in genet detection in managed areas where on-tree lianas were rare.

Samples were first stored at 4 °C and then frozen at −30 °C until DNA extraction. DNA was extracted using the modified 2× CTAB protocol (Murray and Thompson, 1980). Genotypes were determined through PCR amplification of ten microsatellite loci described in Mori et al. (2016b). PCR products were analyzed on a 3130 Genetic Analyzer (Applied Biosystems, CA, USA). Electropherograms were checked for peak patterns using GeneMarker v1.95 (https://softgenetics.com/products/genemarker/).

### 2.5 Data Analysis

#### 2.5.1 Identification of clones

Clones were identified based on the methods proposed by Arnaud-Haond et al. (2007). The ability of the microsatellite markers to distinguish multilocus genotypes (MLGs) was tested by calculating the number of MLGs for all combinations of a given locus, and the results were then verified based on the plateaus of the genotype accumulation curve (Fig. S7). To ascertain whether stems of the same MLG belonged to the same clone, the probability of a given MLG occurring in a population under Hardy-Weinberg equilibrium was calculated using the equation 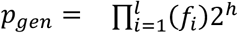(Parks and Werth, 1993), where *f*_*i*_ is the frequency of each allele at the *i*-th locu estimated with a round-robin method, and *h* is the number of heterozygous loci. Then, the probability of obtaining *n* repeated MLGs from a population more than once by chance in N samples (*p*_*sex*_) was calculated using the equation 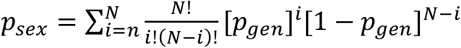(Parks and Werth, 1993). Both *p*_*gen*_ and *p*_*sex*_ were calculated using the R package “RClone” (Bailleul et al., 2016). To distinguish each distinct MLG that belonged to a distinct clone, multilocus lineages (MLLs) were defined based on pairwise genetic distances that have been genotyped in this study (Fig. S8). This procedure was necessary to prevent the false detection of clones due to slightly different MLGs resulting from somatic mutation or genotyping errors. The pairwise genetic distance threshold was determined with the cutoff_predictor function of the R package “poppr” (Kamvar et al., 2014; Kamvar et al., 2015). MLLs are equivalent to genets (clones) in ecological studies, and we will use “genets” hereafter for MLLs. We use “single-ramet genet” for a genet represented by one ramet and “multiple-ramet genet” for a genet represented by two or more ramets.

### 2.5.2 Quantifying clonal contribution and genet areal metrics

We quantified (i) the proportion of clonal ramets, (ii) the number of genets and their ramet-size distribution, and (iii) genet areal metrics. The proportion of clonal ramets was defined as 1 − (number of ramets in single-ramet genets / total number of ramets) following Mori and Kamijo (2025). For each genet, we constructed a minimum convex polygon (convex hull) from ramet coordinates in QGIS 3.30 (QGIS Development Team, 2024). Cumulative genet coverage was calculated as the area of the union of all genet hull polygons within the 21.6-ha study area, with overlapping regions counted only once; we also computed coverage restricted to on-tree ramets to exclude pre-climbing individuals and to provide a concise reference for the post-climbing stage, which is typically the focus of liana ecology and management.

The ramet-size distribution (ramets per genet) was summarized using predefined categories (1, 2, 3, 4–5, 6–10, 11–20, >20) set a priori to preserve resolution where most genets occur and to aggregate the long tail. The genet-area distribution was visualized using an empirical cumulative distribution function (ECDF) for genets with ≥ 3 ramets; in the ECDF, the y-axis gives the cumulative proportion of genets whose area is at or below the corresponding value on the x-axis, and the x-axis (area in ha) was plotted on a logarithmic scale.

### 2.5.3 Genet distribution relative to pre-fragmentation forest

To assess whether observed patterns are consistent with clonal expansion prior to fragmentation or instead with expansion occurring only after fragmentation, we evaluated two aspects of genet distribution: (i) whether genets occur across multiple current forest patches, and (ii) whether such genets are disproportionately located within areas that were forest prior to fragmentation. Under a post-fragmentation-only scenario, all genets would be expected to occur within a single current forest patch; in contrast, the presence of genets spanning multiple patches—especially if their ramets are concentrated within pre-fragmentation forest—would be consistent with expansion initiated prior to fragmentation. As the pre-fragmentation baseline, we used the 1959 forest area, the earliest year in our imagery series and the most contiguous (Figs. 2, S4). For each genet, we assigned each ramet to a current forest patch (Fig. S3) and classified the genet as single-patch if all ramets fell within the same patch, or multi-patch otherwise. We then labeled each ramet as inside or outside the 1959 forest area and compared frequencies between ramets belonging to multi-patch genets and those belonging to single-patch genets using Fisher’s exact test. If the proportion within the 1959 forest area was significantly higher for multi-patch ramets, we interpreted this as a pattern consistent with expansion initiated prior to fragmentation. All analyses were performed in R 4.3.2 (R Core Team, 2023).

## 3 Results

### 3.1 Liana abundance in the fragmented forest

Across the 0.6-ha inventory (15 grid cells), we recorded 9 liana species, totaling 573 stems and a basal area of 0.287 m^2^ (Table S1). *Wisteria floribunda* was dominant with 393 stems, accounting for 69% of total liana stems, and 0.258 m^2^, accounting for 90% of total liana basal area. The next most abundant species by stems was *Parthenocissus tricuspidata* (Vitaceae; 75 stems, 13% of total liana stems), and by basal area was *Toxicodendron orientale* subsp. *orientale* (Anacardiaceae; 0.012 m^2^, 4% of total liana basal area). Together, these values indicate that within the liana community of the fragmented forest, *W. floribunda* is the clearly dominant species by both stem density and basal area.s

### 3.2 Clonal structure

We genotyped 273 *Wisteria floribunda* individuals (231 on-tree, 42 on-floor) and resolved 47 genets, comprising 31 multiple-ramet genets and 16 single-ramet genets (Fig. 3). The proportion of clonal ramets was 94% (257/273). Stage-specific proportions were 94% for on-tree (217/231) and 95% for on-floor (40/42), with no significant difference (Fisher’s exact test; odds ratio = 1.29, 95% CI: 0.28–12.12, P = 1.00). The probability of the study species obtaining a given genotype (*p*_*gen*_ < 0.001) or obtaining repeated genotypes that originated from distinct sexual reproductive events by chance (*p*_*sex*_ < 0.001) were low, indicating that errors in clone identification are unlikely and supporting the robustness of clone assignment.

**Figure 3.**
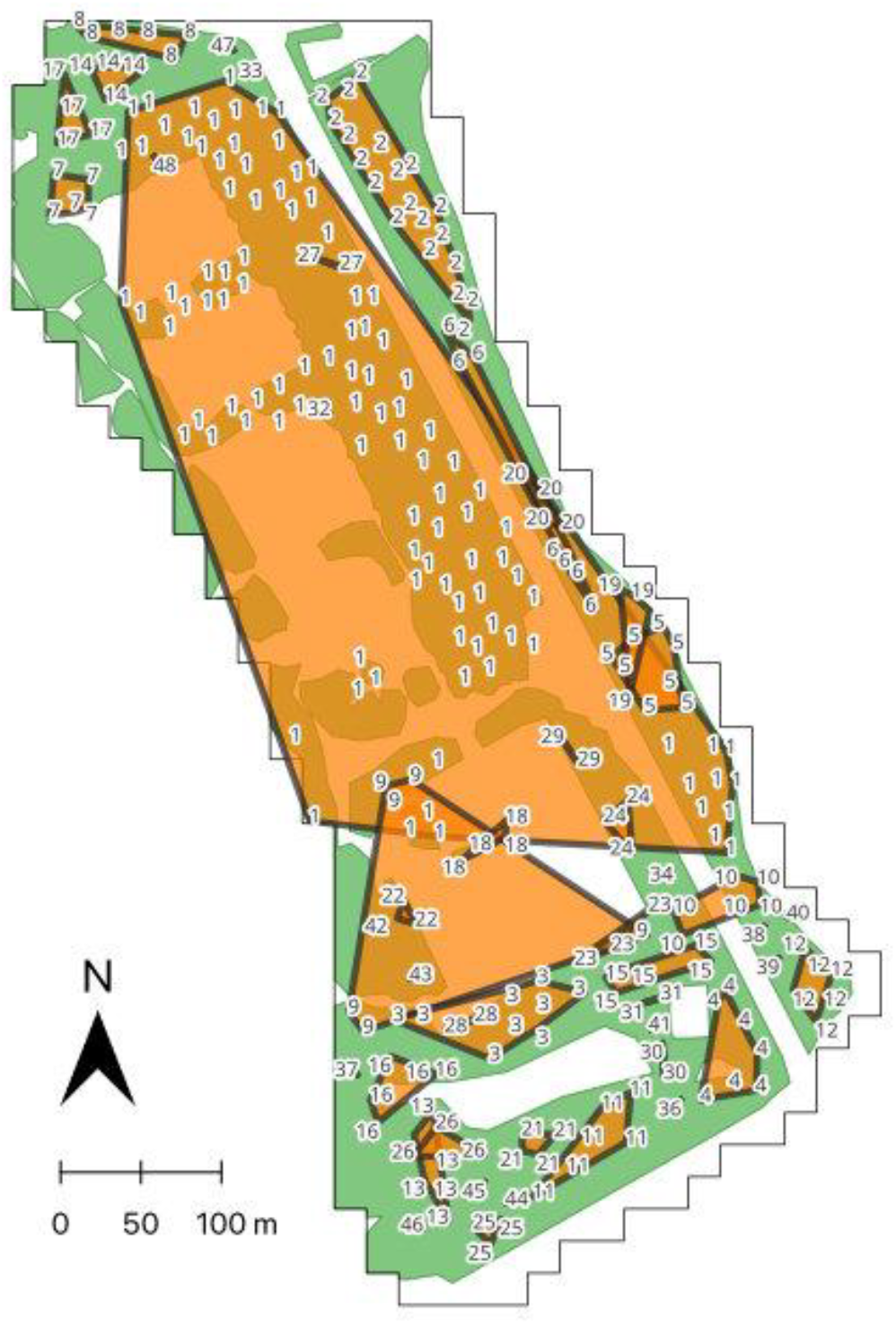
Clonal structure of the study species in the study site. Each number indicate ramets; same numbers indicate genetically identical individuals (i.e., genets/clones). Orange polygons with solid lines indicate genet area obtained as convex hull polygons. Filled green background represent area currently covered by forest.

### 3.3 Genet areal metrics and cumulative coverage

The cumulative genet coverage (union of convex-hull polygons) was 12.5 ha, corresponding to 58% of the 21.6-ha study area. When restricted to on-tree ramets only, cumulative coverage was 12.3 ha (57%), yielding a similar estimate. Among genets with ≥3 ramets (n = 26), genet areas were strongly right-skewed, with a median of 0.060 ha and a range of 0.007–9.5 ha (Fig. 4). The largest genet (ID 1; Fig. 3) comprised 105 ramets (39% of all ramets) and covered 9.5 ha, accounting for 76% of cumulative coverage. These metrics are consistent with large-areal genets contributing substantially to overall coverage in the study area.

**Figure 4.**
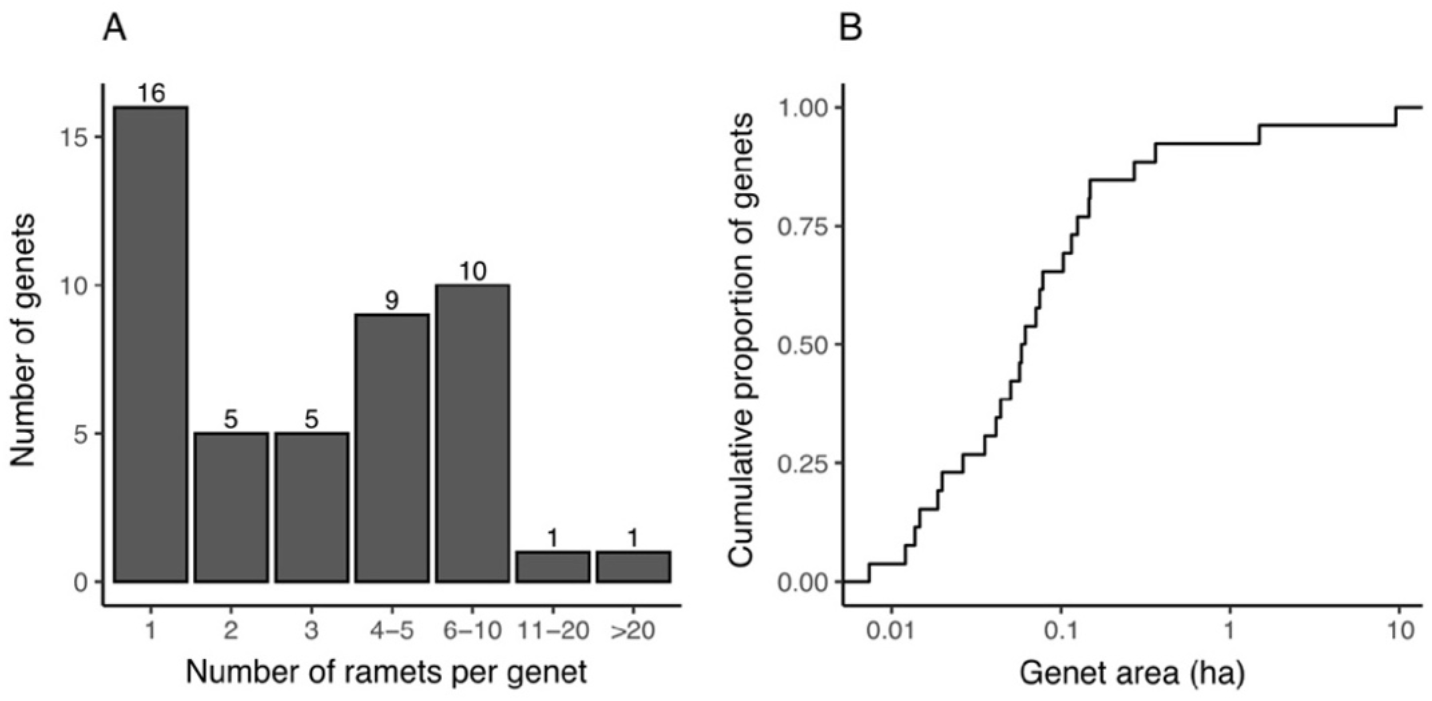
Genet size and area distributions in the fragmented forest. (A) Number of ramets per genet (all genets; n = 47). (B) Empirical cumulative distribution function (ECDF) of genet areas (genets with ≥ 3 ramets; n = 26).

### 3.4 Genet distribution relative to forested areas before fragmentation

Three genets (IDs 1, 9, 10; Fig. 3) occurred across multiple current forest patches (117 ramets; 43% of all ramets), while 44 genets were confined to a single patch (156 ramets). Within areas forested in 1959, 107/117 ramets (91%) from multi-patch genets occurred inside these areas, compared with 95/156 ramets (61%) from single-patch genets. Multi-patch genets had a higher proportion of ramets within areas forested in 1959 than single-patch genets (91% vs 61%; Fisher’s exact test: odds ratio = 6.83, 95% CI: 3.24–15.8, P < 0.001). Taken together, the concentration of multi-patch ramets in forests present in 1959 is consistent with clonal expansion that began before fragmentation, rather than being explained solely by post-fragmentation spread.

## 4. Discussion

This study aimed to identify the demographic source of post-disturbance proliferation in *Wisteria floribunda*—a regionally dominant temperate liana—and to evaluate the spatial scale of that process and its relation to pre-fragmentation forest. Using DNA analysis to estimate the spatial genetic structure across a 21.6-ha fragmented forest landscape, we found that (i) clonality predominated (94% of sampled ramets); (ii) a small number of large genets organized much of the coverage (cumulative 58% of the study area; largest 9.5 ha); and (iii) genets spanning multiple current patches were concentrated within the forest area before fragmentation. Together, these patterns indicate that clonal expansion—rather than seed recruitment alone—accounts for much of the present proliferation across the fragmented forest, and that the present distribution of genets is consistent with expansion initiated prior to fragmentation.

### 4.1 Contribution of clonal expansion to post-disturbance proliferation

Large-scale anthropogenic forest disturbance, including fragmentation, is widely linked to liana proliferation (Laurance et al., 2001; Campbell et al., 2018; Ngute et al., 2024), with edges and gaps creating high-light microsites rich in slender supports where lianas peak in abundance (Putz, 1984; Schnitzer and Carson, 2001; Londré and Schnitzer, 2006). For *W. floribunda* specifically, these conditions are associated with elevated first-year seedling survival (Mori et al., 2020), and seed recruitment would be expected to contribute; nevertheless, our DNA-based assignments indicated that 94% of ramets were clonal. Gap-driven liana increases in tropical forests likewise often include coppiced clonal stems (Ledo and Schnitzer, 2014; Schnitzer et al., 2021). Thus, in this fragmented forest, clonal expansion—not seed recruitment alone—appears to account for most of the present proliferation.

Stage-specific clonal proportions were consistently high for both on-tree and on-floor ramets; because on-floor ramets were sampled only in managed areas lacking on-tree lianas owing to routine mowing, this pattern is compatible with clonality persisting under repeated cutting and with seed-origin cohorts being less represented in such areas. In many systems, liana cutting is followed by resprouting (coppicing) from stem bases and surviving belowground modules, and severed stems can re-establish after cutting (e.g., Alvira et al., 2004; Jarvis-Lowry et al., 2024), which together support the plausibility that rapid post-cutting re-establishment contributes to the very high clonality observed here. More generally, greenhouse experiments show that liana roots can reach fertilized patches faster than tree roots (Putz, 2023), indicating rapid belowground resource foraging that may complement clonal spread after disturbance.

### 4.2 Large genets and pre-fragmentation legacy

Landscape-scale genetic assignments showed a strongly right-skewed distribution of genet areas: a few large genets accounted for much of *W. floribunda* coverage across the fragmented forest.

Comparable size skew and very large, long-lived clones are well documented in clonal plants (Suyama et al., 2000; de Witte and Stöcklin, 2010; Arnaud-Haond et al., 2012). This pattern indicates that post-disturbance proliferation reflects not only widespread clonality but the disproportionate expansion or persistence of a few genets operating over hundreds of meters and across multiple patches. The concentration of coverage in few genets suggests that, in other regions or species, post-disturbance liana increases may likewise be driven by a small number of clones with potentially long-lasting demographic effects.

In addition to the size distribution, the concentration of genets within areas forested in 1959 indicates a legacy component. Genets that span multiple current patches are best explained by legacy genets established before fragmentation that have persisted and/or expanded across areas now delineated as separate patches. Clonal integration among ramets (physiological resource translocation along stolons) offers a plausible mechanism aiding persistence and large areal coverage by supporting re-establishment after partial damage and occupation of favorable microsites (Fahrig et al., 1994; Winkler and Fischer, 2001; Saitoh et al., 2006). Across the population—particularly for genets confined to a single current patch—two pathways likely operate: (i) persistence and expansion of legacy genets and (ii) post-disturbance seed recruitment followed by clonal growth, with pathway (ii) documented for the liana *Trachelospermum asiaticum var. asiaticum* in early-successional forests (Mori and Kamijo, 2025). These lines of evidence indicate that interpreting post-disturbance liana increases requires attention to both pre-fragmentation genetic structure and post-disturbance clonal proliferation, as also suggested by legacy-effect frameworks for forest landscapes (Foster et al., 2003; Johnstone et al., 2016).

### 4.3 Implications for management and monitoring

Lianas are difficult to manage in fragmented forest landscapes, where their abundance is typically high and commonly attributed to high-light conditions and plentiful slender host supports

(Campbell et al., 2015; Campbell et al., 2018). Liana cutting is widely applied in forestry and restoration (Marshall et al., 2020) and has been proposed as a cost-effective natural climate solution that can increase aboveground carbon sequestration and timber yields (Putz et al., 2023). Nevertheless, our findings—multi-hectare genets and a predominance of clonal ramets—indicate that management and control in fragmented forests are especially challenging because post-disturbance proliferation reflects not only local resprouting but also landscape-scale clonal expansion. Consistent with this interpretation, resprouting and re-establishment after cutting or mowing are documented for several lianas (e.g., Alvira et al., 2004; Kartzinel et al., 2015; Jarvis- Lowry et al., 2024). Accordingly, assessment, monitoring, and management may benefit from a genet-based perspective—complementing ramet counts with DNA-based genet assignment and coverage mapping to identify large genets that disproportionately contribute to proliferation, and to help prioritize and evaluate cutting/mowing interventions in fragmented forests.

Clonal capacity varies across species and environments. For example, in a 6-ha plot of closed-canopy old-growth forest (Masaki et al., 1992; Nakashizuka and Matsumoto, 2002), four co-occurring liana species showed marked interspecific contrasts in clonal structure, with *W*. floribunda exhibiting the strongest dependence on clonality and becoming dominant within the liana community (Mori et al., 2021). As contextual reference, values from the same forest stand (Fig. S9; Mori et al., 2018) indicate lower clonal contribution and coverage than observed in the present fragmented forest (e.g., clonal proportion of on-tree ramets 45% vs 94%; cumulative genet coverage 7% vs 57%). Together, these considerations indicate that effective monitoring and management should be species-specific, explicitly consider species’ life histories and disturbance histories, and, where feasible, be grounded in genet-level understanding.

### 5. Conclusions

Landscape-scale, DNA-based genet–ramet assignments indicate that proliferation of *W*.

*floribunda* in this fragmented forest is predominantly clonal, that coverage is organized by a small number of large genets, and that the concentration of multi-patch genets within areas forested in 1959 is consistent with a pre-fragmentation legacy component. Together, these lines of evidence suggest that post-disturbance increases can be accounted for largely by clonal expansion, with seed recruitment playing a more limited role in the present system.

Conceptually, these findings underscore the value of making clonality explicit—both the presence/absence of clonality and the spatial scale of genets—when interpreting liana responses to large-scale anthropogenic forest disturbance. For practice, monitoring and interpretation are likely to benefit from a genet-based perspective (e.g., genet coverage and occurrence across patches), supported where feasible by genetic analysis, alongside conventional above-ground observations, and from approaches that are species-specific and attentive to life histories and disturbance history.

The scope of inference is necessarily limited: evidence comes from a single species in a single landscape. Future work that replicates across species and landscapes, and links spatial distribution of genets with long-term demographic monitoring will clarify when seed establishment versus clonal expansion sustains liana increases and will refine predictions for fragmented forests.

## Acknowledgements

This study was supported by JSPS KAKENHI (grant numbers 21K14882; 24K01802).

## Appendix A. Supplementary material

## Data availability

Data will be made available on request.

