## Supplementary figures and images for "Clonal expansion accounts for most post-disturbance proliferation of a dominant temperate liana, *Wisteria floribunda*, across a fragmented forest landscape"

### Fig. S1

**Figure S1**. A photograph of a typical forest in the study site.

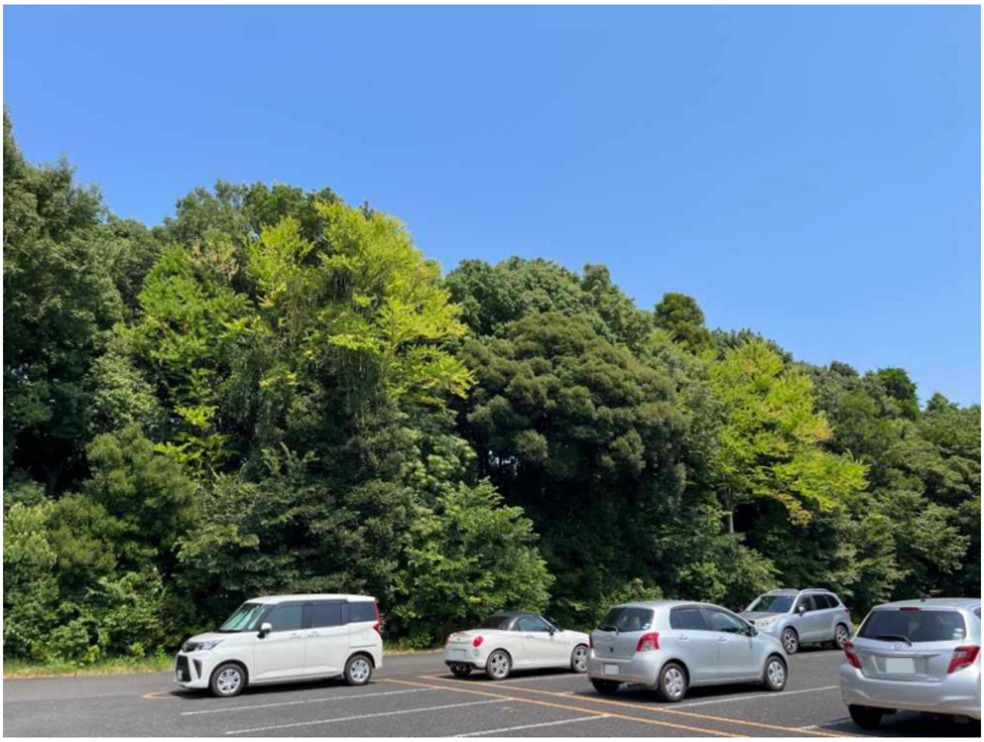

### Fig. S2

**Figure S2.** Temporal changes in forest area and the number of forest patches at the study.

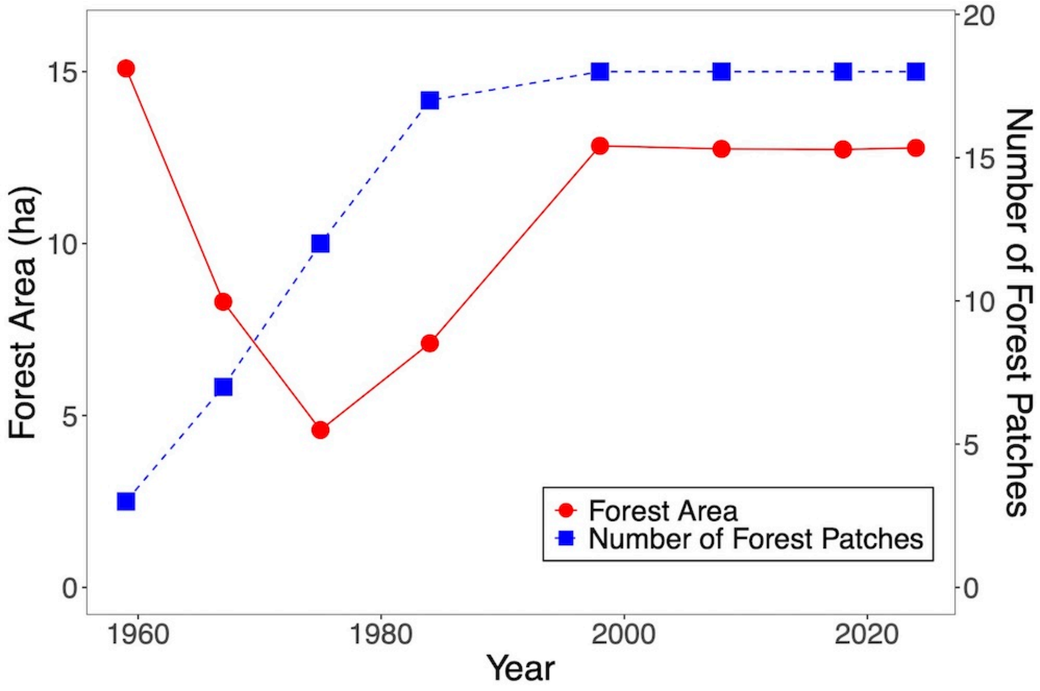
