## Supplementary material for "Clonal expansion accounts for most post-disturbance proliferation of a dominant temperate liana, *Wisteria floribunda*, across a fragmented forest landscape": Fig. S3

**Figure S3.** Fragmented forest patches in the study site. Numbers indicate identifier of fragmented forests with one or more liana stems found in the study site.

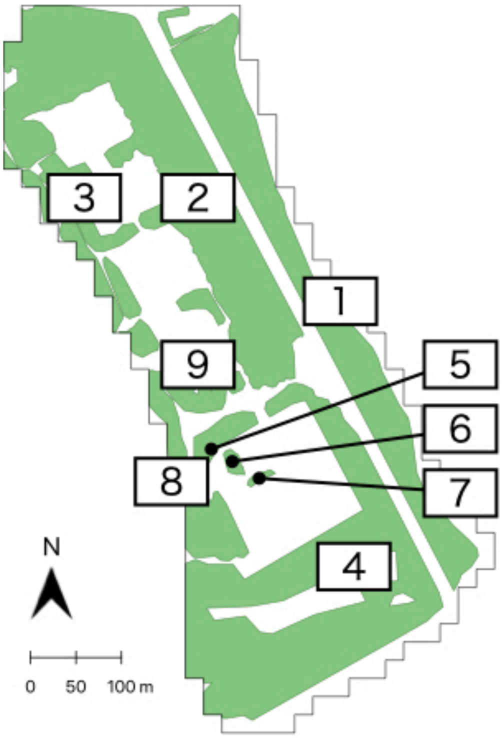
