## Supplementary material for "Clonal expansion accounts for most post-disturbance proliferation of a dominant temperate liana, *Wisteria floribunda*, across a fragmented forest landscape": Fig. S4

**Figure S4.** Managed zones (shaded areas) in the study site. These zones are mowed 2–3 times annually, typically between May and November.

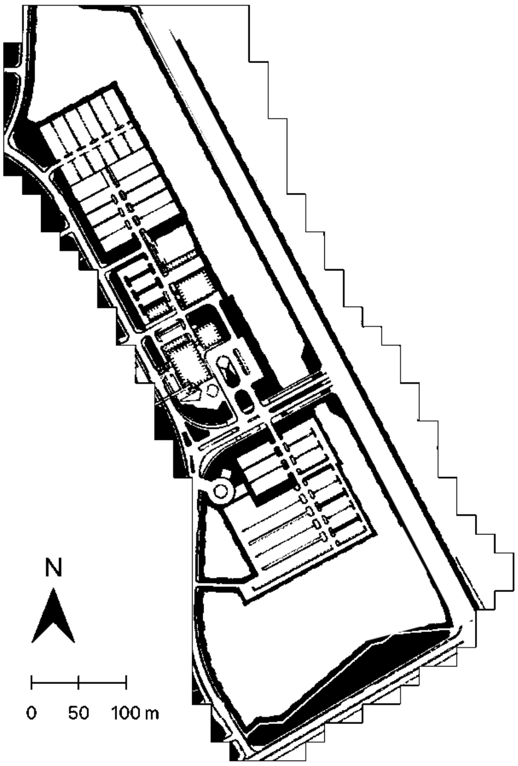
