## Supplementary material for "Clonal expansion accounts for most post-disturbance proliferation of a dominant temperate liana, *Wisteria floribunda*, across a fragmented forest landscape": Fig. S5

**Figure S5.** Photographs of *Wisteria floribunda* (a) at the on-tree stage and (b) the on-floor stage in the fragmented forest at the study site. (c) *Wisteria floribunda* flowering within the study site in early spring.

(a) On-tree individual

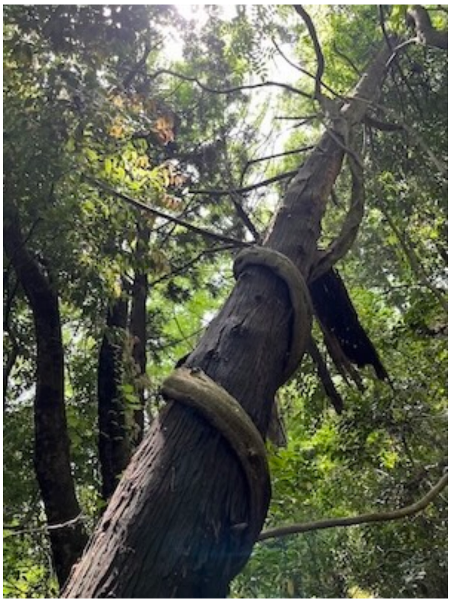

(b) On-floor individual

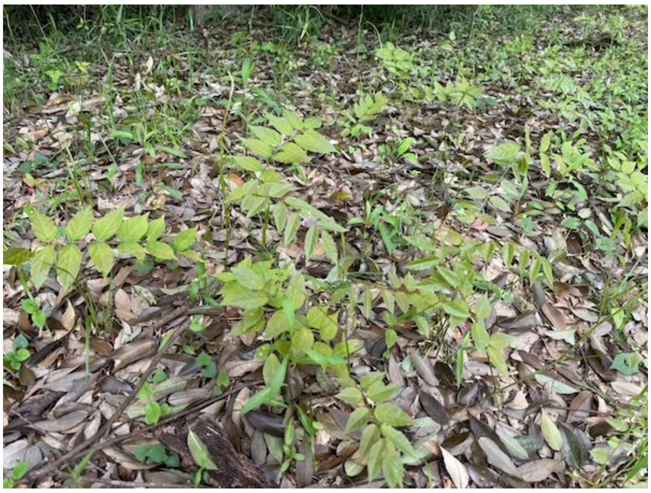

(c) The study species along the forest edge

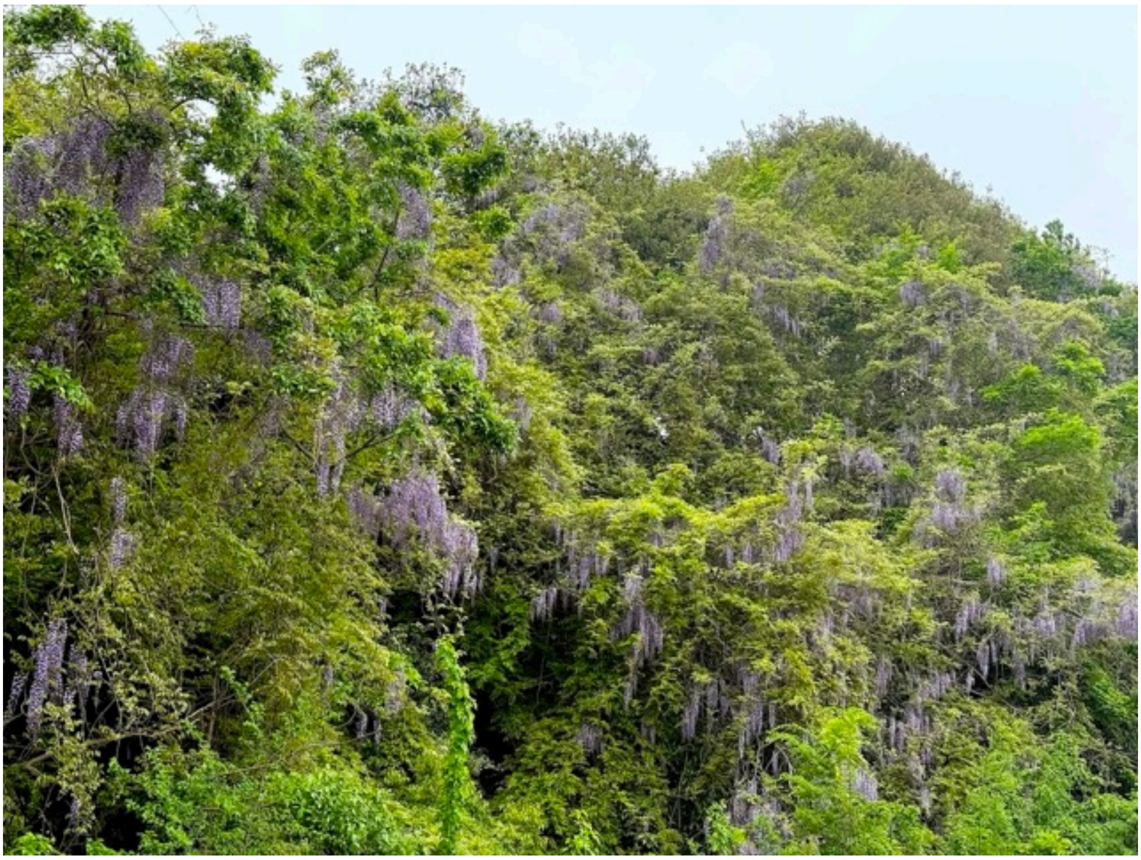
