## Supplementary material for "Clonal expansion accounts for most post-disturbance proliferation of a dominant temperate liana, *Wisteria floribunda*, across a fragmented forest landscape": Fig. S6

**Figure S6.** 20 m x 20 m grid cells utilized in this study. Red grid cells (N = 15) indicate locations of liana inventory conducted in this study.

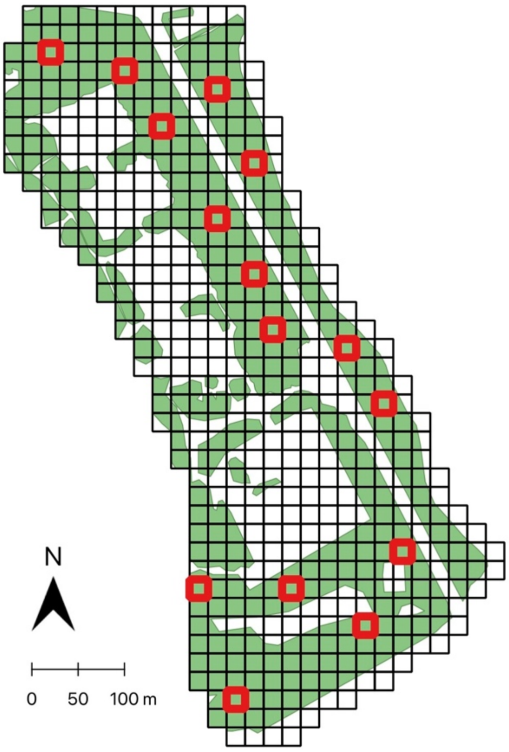
