## Supplementary material for "Clonal expansion accounts for most post-disturbance proliferation of a dominant temperate liana, *Wisteria floribunda*, across a fragmented forest landscape": Fig. S7

**Figure S7.** Genotype accumulation curve. Horizontal dashed lines indicate the total number of multilocus genotypes identified using all SSR loci.

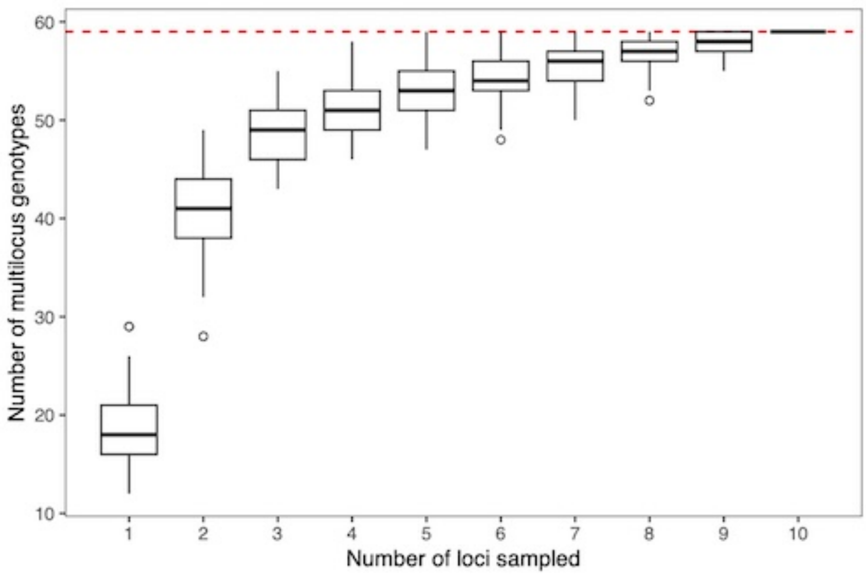
