## Supplementary material for "Clonal expansion accounts for most post-disturbance proliferation of a dominant temperate liana, *Wisteria floribunda*, across a fragmented forest landscape": Fig. S8

**Figure S8.** Pairwise genetic distance of collected samples of the study species. Red vertical line indicates estimated threshold for the genetic distance cutoff.

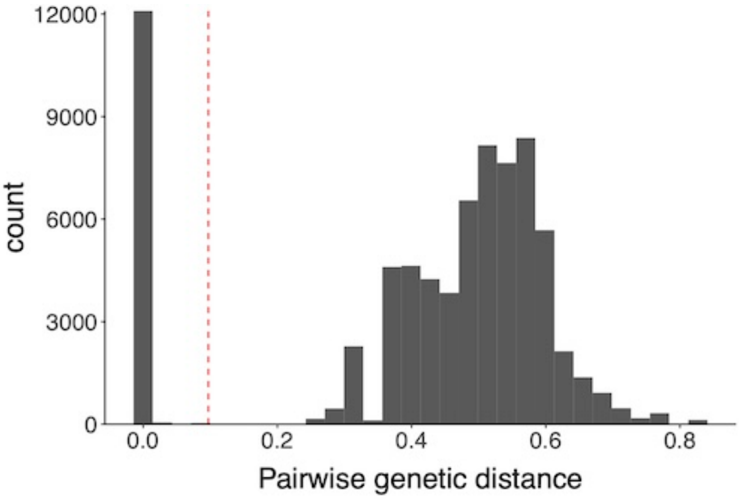
