## Supplementary material for "Clonal expansion accounts for most post-disturbance proliferation of a dominant temperate liana, *Wisteria floribunda*, across a fragmented forest landscape": Fig. S9

**Figure S9**. Clonality (upper: clonal structure maps, lower: clonality metrics) of the study species in the present study site (left column) and the continuous old-growth forest site (right column). Maps are shown in the same spatial scale. For visibility purposes, clonal ramets distributed within the genet polygons are not shown in this figure. Metrics for clonality of the continuous old-growth forest site was calculated based on the same methodology of the present study by sub-sampling ramets in 20 x 20 m quadrats with 1,000 iterations, and mean values and 95% CIs are shown. Clonality of the continuous forest site is based on Mori et al. (2018).

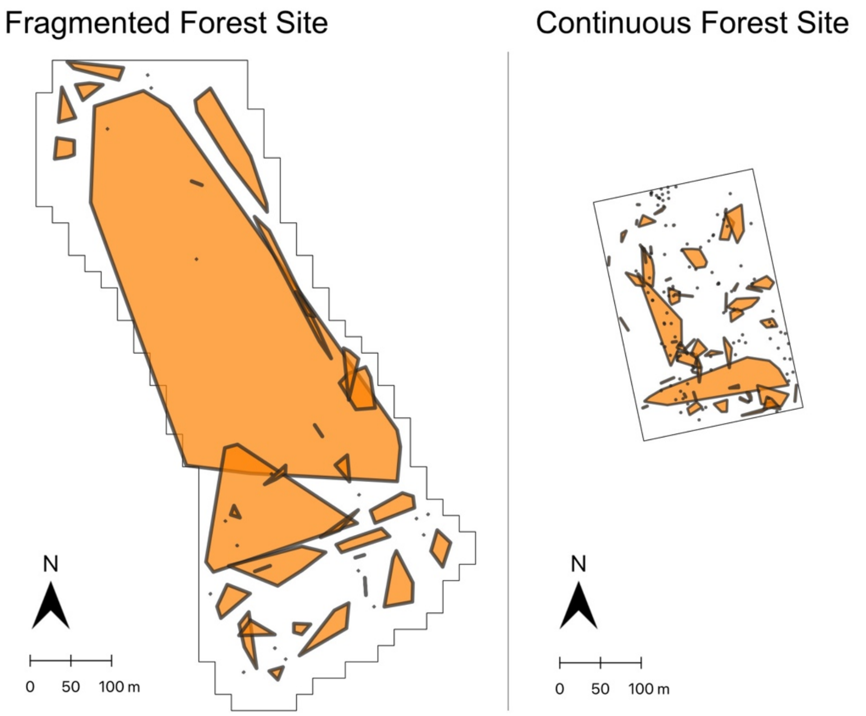


| Metric | Fragmented Forest Site | Continuous Forest Site |
|---|---|---|
| % Clonal ramets | 94% | 45% (38–52%) |
| % Genet area | 57% | 7% (4–10%) |
