## Supplementary material for "Clonal expansion accounts for most post-disturbance proliferation of a dominant temperate liana, *Wisteria floribunda*, across a fragmented forest landscape": Table S1

**Table1**

**Table 1.** Summary of liana community structure investigated in 15 quadrats (total 0.6 ha) located in the fragmented forests in the present study. Species are ordered by stem density. Nomenclature follows Yonekura and Kajita (2008).

| **Family** | **Species** | **Stem Density** | | | **Basal Area** | **Stem Diameter (mm)** | | | |
|---|---|---|---|---|---|---|---|---|---|
| | | **Total** | **Mean** | **SD** | **($m^2$ 0.6 $ha^{-1}$)** | **Mean** | **SD** | **Min** | **Max** |
| Fabaceae | *Wisteria floribunda* | 393 | 26.2 | 17.0 | 0.258 | 18.7 | 22.1 | 1.5 | 144.6 |
| Vitaceae | *Parthenocissus tricuspidata* | 75 | 6.2 | 6.6 | 0.008 | 7.7 | 8.5 | 1.0 | 47.9 |
| Lardizabalaceae | *Akebia trifoliata* | 61 | 5.5 | 5.5 | 0.006 | 9.8 | 6.0 | 2.4 | 28.1 |
| Lardizabalaceae | *Akebia quinata* | 12 | 4.0 | 4.4 | < 0.001 | 4.7 | 3.1 | 1.3 | 12.5 |
| Araliaceae | *Hedera rhombea* | 12 | 4.0 | 3.6 | 0.002 | 7.5 | 10.7 | 1.2 | 30.5 |

| | | | | | | | | | |
|---|---|---|---|---|---|---|---|---|---|
| Anacardiaceae | *Toxicodendron orientale* subsp. *orientale* | 7 | 1.8 | 0.5 | 0.012 | 36.1 | 31.2 | 2.5 | 80.9 |
| Fabaceae | *Pueraria lobata* subsp. *lobata* | 6 | 6.0 | | < 0.001 | 11.0 | 8.6 | 1.5 | 19.0 |
| Apocynaceae | *Trachelospermum asiaticum* var. *asiaticum* | 6 | 3.0 | 1.4 | < 0.001 | 8.1 | 6.8 | 1.2 | 16.1 |
| Caprifoliaceae | *Lonicera japonica* | 1 | 1.0 | | < 0.001 | 1.4 | | 1.4 | 1.4 |
| Total | | 573 | 38.2 | 21.5 | 0.287 | 15.8 | 19.7 | 1.0 | 144.6 |
